# Rescue of tomato yellow leaf curl virus mutants with heterologous iterons through *in planta* evolution

**DOI:** 10.1101/2025.05.27.656274

**Authors:** Khwannarin Khemsom, Ruifan Ren, Junping Han, Camila Perdoncini Carvalho, Eric Matthew Snider, Deyong Zhang, Feng Qu

**Author notes:** **Correspondence:** Feng Qu. These authors contributed equally to this study (Khwannarin Khemsom, Ruifan Ren).

## Abstract

The single-stranded, circular DNA genomes of geminiviruses contain iterated motifs of 5-6 nucleotides, known as iterons, upstream of the replication protein (Rep) coding region. Iterons were previously found to interact with cognate Rep in a sequence-specific manner, and the iteron-Rep interaction was needed for viral DNA replication. Nonetheless, iterons of closely related viruses often have different sequences, suggesting diversifying selection. To identify selection pressures driving iteron diversification, we constructed tomato yellow leaf curl virus (TYLCV, isolate SH2) mutants in which the iteron motifs were replaced with those of closely related tobacco curly shoot virus (TbCSV, isolate Y35). All mutants replicated in inoculated leaves of *Nicotiana benthamiana*, but many failed to spread systemically. However, the systemic movement defects were mostly rescued by *de novo* mutations. Intriguingly, these *de novo* mutations did not restore the iterons to SH2 sequences. Rather, they likely enabled viral escape from repression exerted by the heterogenous Y35 iterons absent of a matching Rep. These results are consistent with iterons acting as sites of competitive binding by host-encoded transcription factors (TFs) and the cognate Rep. The iteron-TF binding commences as soon as viral genomes enter cell nuclei, committing genome copies to Rep mRNA transcription and protein translation; but also blocking them from replication. Conversely, iteron-Rep binding is possible only after Rep is produced, and likely repels TFs from some genome copies, permitting replication initiation. Testing this model through future research should clarify the intricate evolutionary interplays between geminiviruses and their crop hosts, and inform novel management strategies.

**Author Summary:** Geminiviruses are important crop pathogens worldwide for which effective control measures are lacking, due to incomplete understanding of their evolutionary dynamics in infected plants. The current study focuses on a class of short sequence repeats in geminiviral genomic DNA, known as iterons, sitting immediately upstream of the viral gene encoding replication protein (Rep). Iterons are interesting because even though their positions and repeat patterns are conserved across all geminiviruses, their sequence identities are highly diverse. Our investigations revealed that contrary to previous reports, the sequence identity of iterons is non-essential for tomato yellow leaf curl virus (TYLCV) to replicate. Rather, they are repressors of replication, and this repression is overcome by their binding with cognate Rep. Our findings led to a new model postulating that the genome section encompassing iterons likely evolved specific sequence motifs to entice host-encoded transcription factors (TFs), facilitating rapid Rep production. Conversely, Rep promotes viral replication by removing TFs from genome copies through competitive iteron binding. Future testing of this new model will likely unveil novel targets for more effective management of crop diseases caused by geminiviruses.

## Introduction

Viruses of the family *Geminiviridae* are among the most common pathogens of crop plants (1), causing devastating losses in staple crops such as cassava, cotton, maize, soybean, tomato, and wheat. These viruses have relatively small, single-stranded (ss), circular DNA genomes, encoding up to a dozen viral proteins (2). Geminiviruses replicate their genomes in host cell nuclei by recruiting a host-encoded DNA-dependent DNA polymerase (DNA Pol). Nevertheless, they do encode an accessory replication protein, designated variously as Rep, C1, AC1, or AL1 depending on viruses under investigation, that recruits DNA Pol to geminiviral DNA. The Rep protein is also responsible for specifying the rolling circle replication mode, enabling a single copy of a circular genome [designated as a (+)-strand] to template the synthesis of multiple copies of complementary (-) strand intermediates. Upon circularization, these (-)-strand intermediates each template the synthesis of numerous copies of progeny (+) genomes, ensuring productive virus reproduction.

In addition to encoding various proteins, the circular ssDNA genomes of geminiviruses also harbor various *cis*-acting elements. The most important among them is a DNA stem-loop consisting of a double-stranded stem of 11 base pairs (bp) and a single-stranded loop featuring a highly conserved TAATATTAC motif. Inside this motif lies the site of Rep-mediated nicking of circular genomes (TAATATT/AC), a key step required for the initiation of rolling circle replication (3,4). This DNA stem-loop is indispensable for geminivirus replication, and indeed must be duplicated at both ends of a linear, unit-length genome to ensure infectivity of a geminiviral infectious clone (5) (also see Fig. 1A).

**Figure 1.**
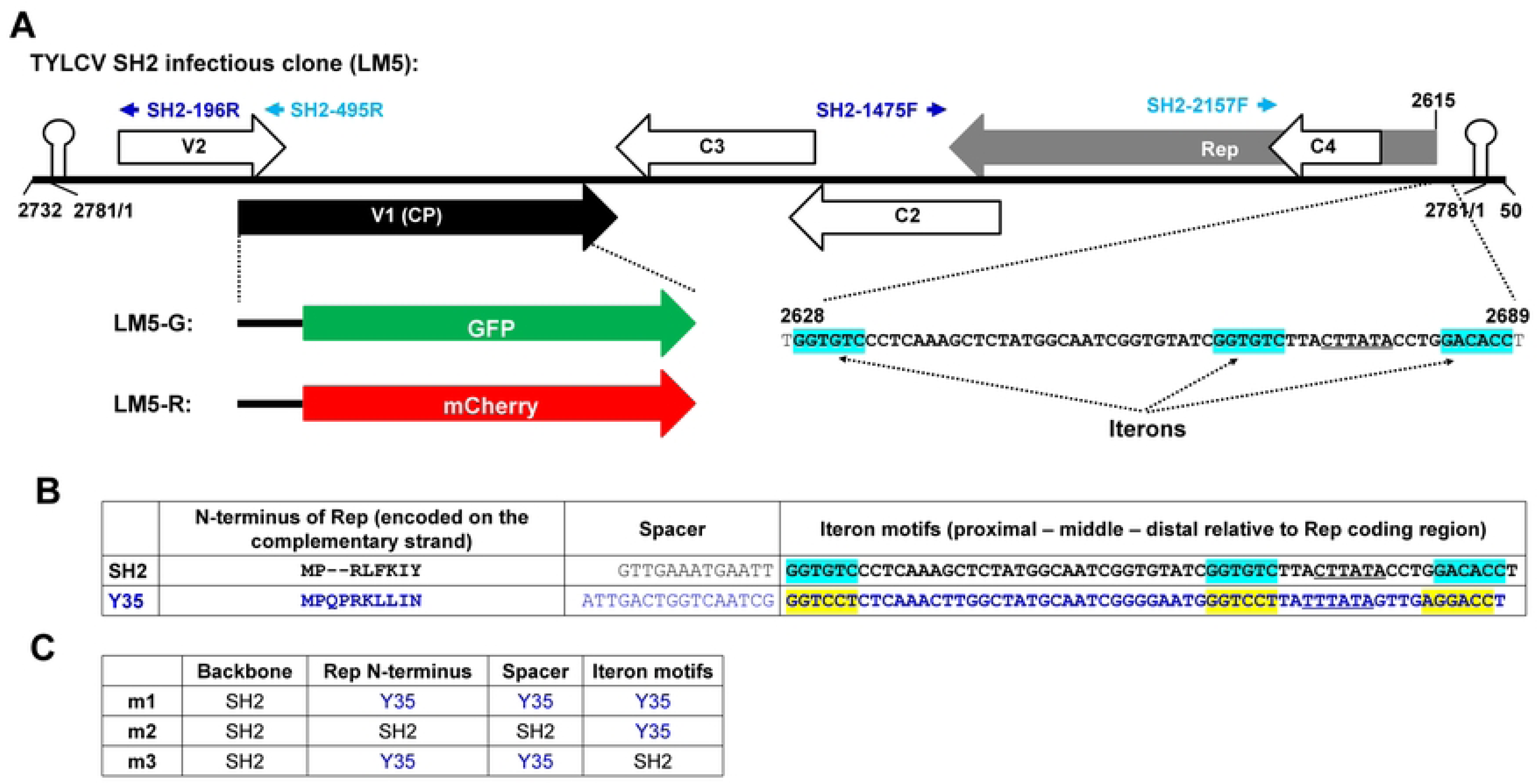
**A**. Schematic representation of the TYLCV SH2 infectious clone LM5, and its two derivatives LM5-G and LM5-R. The linearized form of the full-length SH2 genome, plus 50-nt duplications at both ends, were inserted in the binary plasmid pAI101 (50). The large arrows with internal labels of Rep, C4, C2, C3, V1, and V2 denote TYLCV-encoded proteins. In LM5-G and -R, the V1 coding region was modified to accommodate GFP and mCherry inserts, respectively. Note that LM5-G and -R differ from LM4-G and -R reported earlier (16) – the N-terminal nuclear localization signal (NLS) of V1 has now been decoupled from GFP and mCherry in these new constructs. The small light and blue arrows on the top denote two pairs of primers used for specific detection of circularized form of the replication-generated viral DNA. Finally, the 62-nt from positions 2628-2689 encompass the 3 SH2 iterons, with the specific iteron motifs painted light blue. **B**. Comparing the SH2 and Y35 iterons, spacers between iterons and Rep coding sequences, and the first 8/10 amino acid residues of Reps. Y35 nt and aa are in blue fonts, and the Y35 iterons are painted yellow. Note the Reps are coded by the (-) strands of the respective viral genomes. **C**. Compositions of the three chimeric mutants, m1, m2, and m3.

A second class of *cis*-acting motifs, referred to as iterons, are short (5-6 nucleotides or nt) sequence repeats located upstream of the Rep protein coding region (6,7). Iterons typically repeat for 3-4 times, with at least one of the repeats being antisense to others (7–10). Iterons have been shown to mediate sequence-specific binding with Rep protein of the same virus (11–14), and this iteron-Rep interaction was found to be needed for successful viral replication. Intriguingly, the iteron-Rep binding required the iteron-containing DNA to be double-stranded (10,13), suggesting that geminiviral genomes must either be converted into double-stranded forms, or engage in intra-molecular base-pairing, in order to avail iteron motifs to Rep-binding.

Although iterons are usually conserved among isolates of the same geminivirus species, they are highly divergent among different, though still closely related virus species. Listed in Supplementary Fig. 1A are 4 examples of closely related geminiviruses/isolates, namely tomato yellow leaf curl virus (TYLCV) isolate SH2, tobacco curly shoot virus (TbCSV) isolates Y41 and Y35, and tomato yellow leaf curl Sardina virus (TYLCSV). These viruses share whole genome nt level identities of more than 75% (Suppl Fig 1B, percentages in black fonts) and Rep protein amino acid (aa) level similarities of more than 87% (Suppl Fig. 1B, percentages in green fonts). In particular, TbCSV Y41 and Y35 are more than 96% identical at both the whole genome nt level and the Rep aa level. Nevertheless, Y41 and SH2, but not Y41 and Y35, share the same, though differently spaced, iteron sequences (8,10) (Suppl Fig. 1C, the iterons are painted light blue and yellow, respectively). Moreover, iterons of TYLCSV have entirely different sequences (Suppl Fig. 1C, iterons are painted purple).

Thus, it appears that iterons, despite their involvement in sequence-specific interactions with cognate Reps, are under rapid diversifying selection. Consistent with this view, Reps, or more specifically the iteron-interacting domains of Reps, also appear to evolve rapidly to maintain the sequence-specific interactions (6,7,9). Indeed comprehensive bioinformatic analyses of large numbers of geminiviruses and nanoviruses identified two separate Rep domains whose aa sequences co-varied with iteron sequences (7,9) (Suppl Fig. 1D). However, it has not been thoroughly examined as to exactly what drives the rapid diversification of iteron motifs and the co-varying iteron-interacting Rep domains.

Here we report an attempt to identify such selection pressures, using TYLCV SH2 as the model virus, and *Nicotiana benthamiana* as the model host (15,16). TYLCV is a monopartite member of the genus *Begomovirus*, family *Geminiviridae*. Its circular ssDNA genome of approximately 2,800 nt encodes at least six proteins (2,17,18). The four proteins (Rep, C2-C4) encoded on the (-)-strand of the genome are early expressing, participating in various aspects of viral genome replication, transcriptional activation, and host defense mitigation (Figure 1A) (2,19). In particular, Rep is absolutely required for the rolling circle replication of TYLCV genome (20–22). The two proteins encoded on the (+)-strand of TYLCV genome, known as V1 and V2, are late expressing, and function as capsid protein (CP) and suppressor of RNA silencing, respectively (Figure 1A). V1 (CP) is not essential for replication of TYLCV (23), but needed for the intra- and inter-cellular trafficking of the virus (24).

In the current investigation, we modified the TYLCV SH2 genome by replacing its iterons with those of TbCSV Y35 (Fig. 1). We then infected *N. benthamiana* plants with the modified viruses, and subjected infected plants to systematic analyses for up to 9 weeks. Identification of *de novo* mutations, and subsequent examinations of these new mutations in plants, led us to conclude that, absent of a cognate Rep, iterons act to repress virus replication. Such replicational repression was probably caused by recruitment of host-encoded transcription factors (TFs) by iterons or other tightly linked sequence motifs. The recruited TFs enhanced Rep mRNA transcription to enable rapid accumulation of Rep proteins, but also sequestered viral genome copies from replication. While wildtype TYLCV could ease this sequestration through iteron-Rep binding, some of the viral mutants we examined probably escaped the same sequestration through spontaneous mutations that weakened iteron-TF binding. Such mutation-driven escape may prime the diversification of iteron motifs in closely related geminiviruses.

## Results

### Replacing all iterons of TYLCV SH2 with their TbCSV Y35 counterpart compromises, but does not abolish, viral replication in single cells

Earlier authors found that iterons and Rep of the same geminivirus interacted with each other in a highly sequence-specific manner, and such iteron-Rep interaction was required for replication of viral genomes (6,8,9,11–14,25). However, these studies mostly used viruses with bipartite genomes, or those hosting satellite DNAs, with iteron-disrupting mutations engineered in the non-Rep-encoding genome segments (e.g. DNA-B of bipartite geminiviruses, or satellite DNAs). To determine whether iteron perturbation in *cis* compromises replication of a monopartite geminivirus, we manipulated the genome of TYLCV SH2 to generate three mutants – m1, m2, and m3 (Fig. 1C). In m1, the 97-nt genome section of SH2 encompassing all three iterons, a 13-nt spacer, and the 24 nt encoding the N-terminal 8 aa of SH2 Rep was replaced with the 105-nt counterpart in TbCSV Y35 (Fig. 1B, C). The reason for also exchanging Rep N-terminus was because this domain, along with another Rep domain approximately 60-aa downstream, was found to co-vary with iteron sequences (7,9) (Suppl Fig. 1D). By contrast, in m2 only the 60-nt iteron region was exchanged, whereas in m3 only the 8-aa (24 nt) Rep N-terminus and 13-nt spacer were replaced (Fig. 1B, C. Sections originated from Y35 are highlighted with blue fonts).

These mutations were first introduced into LM5-G and LM5-R (Fig. 1A), two SH2 derivatives in which the V1 coding region was partially replaced by GFP and mCherry coding sequences. LM5-G and LM5-R were fully capable of replicating in single cells, but were unable to move cell-to-cell due to the truncation in V1. The LM5-G and LM5-R-based m1, m2, and m3 mutants were delivered into *N. benthamiana* leaves via agro-infiltration to assess their replication in single cells. As shown in Fig. 2A, panel a, mixture of wildtype LM5-G and LM5-R replicated robustly to produce brightly fluorescent epidermal cells. Note a substantial fraction of cells supported the replication of LM5-G or LM5-R alone (rather than both), reflecting intracellular reproductive bottlenecks revealed through an earlier study (16).

**Figure 2.**
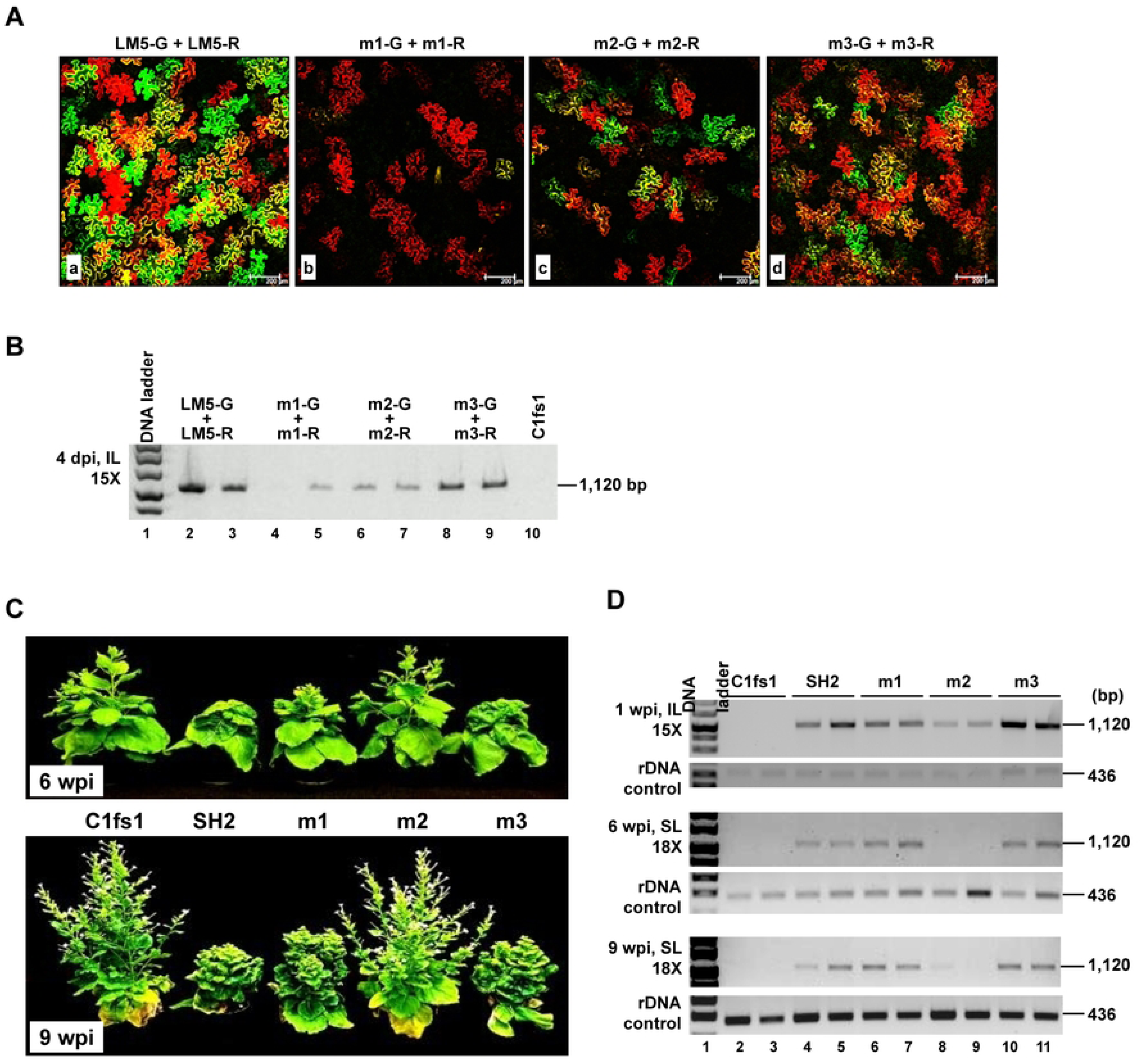
**A**. The m1, m2, and m3 mutants compromises SH2 single-cell replication to varying extents. The m1-G, m1-R, m2-G, m2-R, m3-G, m3-R constructs are derivatives of LM5-G and LM5-R, respectively, with corresponding m1, m2, and m3 mutations. The pairs of constructs were delivered into *N. benthamiana* epidermal cells via agro-infiltration. The cells were observed with a confocal microscope and photographed at 4 days post agro-infiltration. **B.** Semi-quantitative PCR assessing relative accumulation levels of viral genomic DNA in cells receiving constructs shown above the lanes. Note the PCR primers do not distinguish between the GFP- and mCherry-coding constructs. Also note that the non-replicating C1fs1 mutant (containing a frameshift mutation in Rep coding sequence) was included as a negative control. **C**. *N. benthamiana* plants showing symptoms of different SH2 mutants 6 and 9 weeks post inoculation (agro-infiltration) (6 and 9 wpi). **D**. Assessing the relative levels of viral genomic DNA using semi-quantitative PCR in plants treated with m1, m2, and m3 mutants. At 1 wpi, the agro-infiltrated leaves (IL) of two representative plants were analyzed, whereas at 6 and 9 wpi, the systemic leaves (SL) were exmined. The numbers of PCR cycles were also shown. The rDNA control were always amplified for 15 cycles.

With the m1-G & m1-R mix, the replication-dependent GFP and mCherry fluorescence were much weaker, and detectable in fewer cells (Fig. 2A, panel b). The loss of GFP fluorescence was more pronounced so that green-fluorescent cells were barely visible (Fig. 2A, panel b). These observations were also confirmed with the detection of replicating viral DNA with semi-quantitative PCR (Fig. 2B, lanes 4 & 5). Note no PCR fragment of expected size was obtained from the negative control inoculated with the non-replicating C1fs1 mutant (lane 10). Thus, replacing the iterons and Rep N-terminus together severely compromised, but importantly did not abolish, TYLCV replication in single *N. benthamiana* cells.

With m2-G plus m2-R, cells emitting GFP and mCherry fluorescence were fewer, and less bright, than those receiving LM5-G plus LM5-R, but brighter than m1-G plus m1-R (Fig. 2A, compare panels a, b, & c). Consistently, genomic DNA levels of these two mutants were higher than m1-G + m1-R, but lower than LM5-G + LM5-R (Fig. 2B, lanes 2 – 7). Finally, when treated with m3-G and m3-R, cells expressing GFP and/or mCherry were nearly as abundant as those treated with LM5-G and LM5-R, but the fluorescence was slightly less intense. This result was once again verified by semi-quantitative PCR detection of viral genomes. Together these observations demonstrated that these three mutants compromised SH2 replication to varying degrees at the single-cell level.

### Mutants m1 and m3, but not m2, elicit systemic symptoms in *N. benthamiana*

We next migrated the three sets of mutations into the wildtype SH2 infectious clone (LM5, Fig. 1A), and tested the resulting mutants in *N. benthamiana*. The m3-infected plants, similar to SH2-infected plants, began to exhibit systemic symptoms at approximately 2 weeks post inoculation (wpi) (Fig. 2C). The m1-infected plants started to show symptoms by 3 wpi, representing a modest delay of one week. As a result, these plants were slightly taller than SH2-infected plants by 6 wpi (Fig. 2C). By contrast, m2 failed to cause visible diseases in any of the 6 infected plants. The symptom severity differences were consistent with semi-quantitative PCR detection of viral genomic DNA. As shown in Fig. 2D, m2 genomic DNA levels in 1 wpi inoculated leaves (ILs), as reflected by the intensity of PCR products (15 cycles), were lowest among the three mutants (Fig. 2D, top panel, lanes 8, 9). m1 genome levels were higher than m2 but slightly lower than the wildtype SH2 (lanes 4 – 7). Interestingly, m3 genomes appeared to accumulate to levels higher than SH2 in ILs. Sequence analysis of the PCR products verified that all mutations remained stable in ILs at 1 wpi. Notably, when the systemically infected leaves (SLs) were examined at 6 and 9 wpi, both m1 and m3 genomes accumulated to levels indistinguishable from SH2 (Fig. 2D). However, m2 was undetectable at 6 wpi, and was present in just one plant by 9 wpi, at a very low titer. Thus, among the three mutants, m2 was most debilitated at the systemic infection level.

### Both m1 and m2 mutants incur *de novo* mutations in systemically infected leaves

To determine whether the mutations of m1 and m2 were stable in SLs, we next subjected the PCR-amplified genome fragments to sequence analyses. At 6 wpi, viral DNA obtained from 3 m1-infected plants all contained a new mutation at the 4^th^ position of middle iteron, changing the Y35-borne GGTCCT motif to GGT<u>T</u>CT (Fig. 3A, *de novo* mutated nt in red font). Furthermore, one of the 3 DNA samples also contained a 2^nd^ mutation at the 6^th^ position of distal iteron (AGGACC to AGGAC<u>A</u>). Remarkably, neither of the *de novo* mutations returned the iteron motifs to the sequences of wildtype SH2 (GGT<u>T</u>CT vs GGTGTC; AGGAC<u>A</u> vs GACACC; Fig. 1C). Even more interestingly, at 9 wpi the 1^st^ mutation was detected in all 6 plants analyzed, and the 2^nd^ mutation was also stable in the plant in which it first emerged. Also worth noting was that the original nt (a C) at the 4^th^ position of middle iteron was undetectable in any of the samples. These results suggested that robust systemic infection did not require the specific sequences of middle and distal iteron motifs to conform to that of either SH2 or Y35.

**Figure 3.**
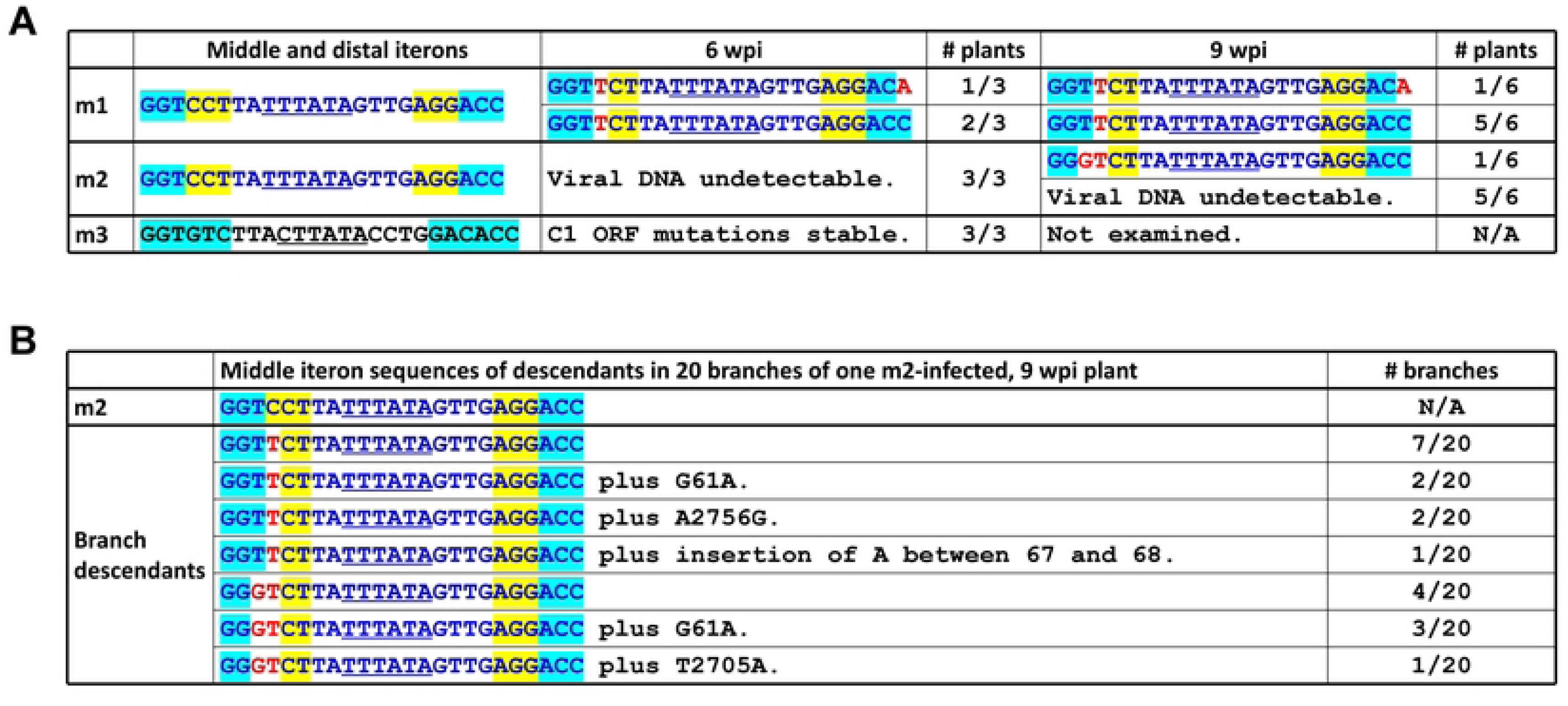
**A & B**. Sequences of m1, m2, and m3 progeny in systemic leaves at 6 and 9 wpi. Only the sequences spanning the middle and distal iterons are shown. Sequences of Y35 origin are in blue fonts, those of SH2 origin are in black fonts. Iteron nts of Y35 origin are painted yellow, those of SH2 origin are painted light blue. The TATA-boxes (in antisense orientation) are underlined. Red nts denote *de novo* mutations emerging from infected plants. The extra mutations occurring in different branches all mapped to the intergenic region between Rep and V2 ORFs.

None of the 3 m2-inoculated plants examined at 6 wpi yielded detectable levels of viral DNA. However, at 9 wpi one of the 6 plants examined contained low levels of viral DNA (Fig. 2D). When subjected to sequence analysis, the PCR-amplified viral DNA contained 2 mutations at 3^rd^ and 4^th^ positions of middle iteron, changing GGTCCT to GG<u>GT</u>CT, deviating the motif sequence even further from that of wildtype SH2 (GGTGTC) or Y35 (GGTCCT). To corroborate this result, we then collected leaf samples from 20 different branches of this plant for DNA extraction and PCR detection of viral DNA. As summarized in Fig. 3B, among the 20 branches, 12 contained descendants that harbored the single C-to-T change also found in m1 descendants, whereas the remaining 8 branches contained variants that had the TC-to-GT change. Together these results showed that when present in SH2 backbone, the Y35-borne middle iteron incurred *de novo* mutations that rendered it dissimilar from both SH2 and Y35.

Finally, progeny of m3 maintained the original m3 mutations, suggesting that the 10-aa N-terminus originating from Y35 was stable in SH2 background, and had minimal impact on viral systemic infections. Together these results demonstrated that when the three iteron motifs were exchanged in concert, the middle motif probably hindered SH2 replication, and its adverse impact must be mitigated through *de novo* mutation(s) in order to restore systemic infection to affected viruses. To reiterate, such new mutations did not convert the motif sequence toward that of SH2 or Y35, thus suggesting a relief from a repressive activity exerted by Y35 iterons in the absence of the cognate Y35 Rep.

### *De novo* mutations bolster infectivity of the m1 mutant

It is worth emphasizing that the original middle iteron of the m1 mutant, of Y35 origin, was never detected in the systemic leaves of plants we examined. We thus speculated that the single C-to-T *de novo* mutation was needed for m1 descendants to spread systemically. To test this, we introduced this mutation back into m1. For comparison, we also introduced the C-to-A distal iteron mutation found in one of the m1 descendants, as well as the TC-to- GT mutations recovered from m2 descendants, into m1. The new mutants are designated as m1a, m1b, and m1f (Fig. 4A, *de novo* mutations highlighted in red font). As shown in Fig. 4B, all three of the new mutants accumulated genomic DNAs to levels similar to that of SH2 and m1 in inoculated leaves. Both m1a and m1f caused symptoms that emerged at about the same time as SH2, thus one week earlier than m1 (Fig. 4C). Curiously, the m1b mutant harboring the TC-to-GT double mutations developed symptoms slightly later than m1a and m1f, causing the infected plants to be slightly taller (Fig. 4C).

**Figure 4.**
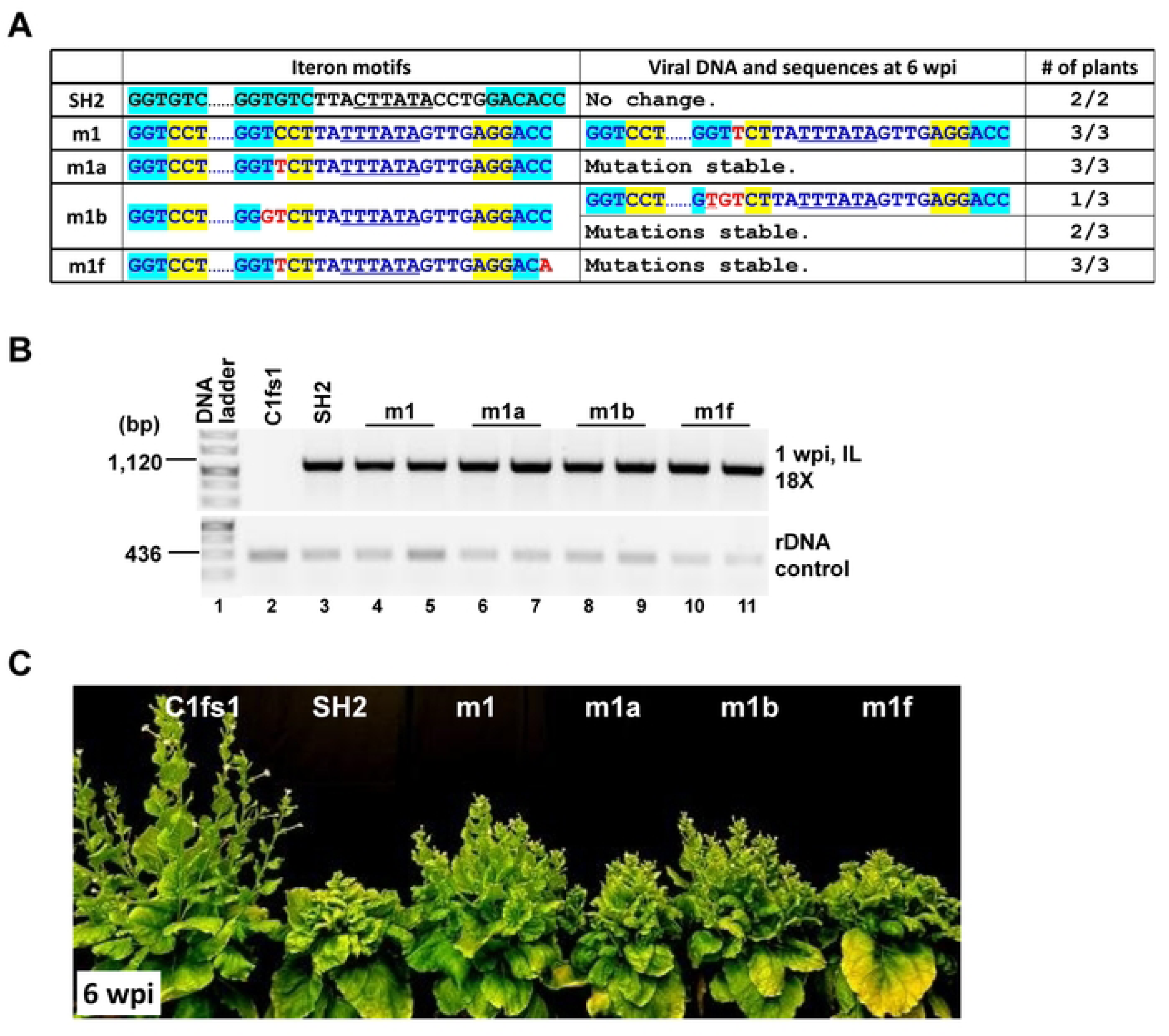
*De novo* mutations identified in m1 and m2 descendants enhance m1 symptoms. **A**. Iteron portion sequences of new m1-based mutants incorporating the *de novo* mutations; and sequences of their descendants at 6 wpi. **B**. Relative accumulation levels of m1, m1a, m1b, and m1f mutants in IL at 1 wpi. **C**. Symptoms of infected plants at 6 wpi.

When subjected to sequence analyses at 6 wpi, the three new m1-infected plants once again yielded viral progeny that incurred the C-to-T *de novo* mutation (Fig. 4A). Combined with the earlier onset of m1a disease symptoms, the highly reproducible emergence of C-to-T mutation strongly suggested that this mutation was responsible for the systemic infections of m1 descendants. Conversely, the original m1 mutant must have replicated poorly, thus easily overtaken by the m1a variant containing C-to-T mutation. Furthermore, this C-to-T mutation, upon emergence in the m1 backbone, was apparently very stable, as it did not evolve further into TC-to-GT. This was unlike the same C-to-T mutation incorporated in the m2 backbone, where it did evolve further into TC-to-GT (see later). Finally, plants infected with the m1f mutant were as severely infected as m1a-infected ones, with both of its mutations remaining stable in plants (Fig. 4B and C). Together these results indicate that the *de novo* mutations detected in m1 descendants correlated with superior infectivity.

### *De novo* mutations restore infectivity to m2 mutant without converting middle iteron to SH2 motif

Unlike the m1-infected plants, those infected with m2 did not have any systemic symptoms, and most of them also failed to accumulate viral genomic DNA (Figs. 2C and 5C). Indeed, even the single plant in which m2 acquired the C-to-T and TC-to-GT mutations did not have visible systemic symptoms. We thus set out to resolve whether these *de novo* mutations also improved the infectivity of the m2 mutant. To this end, the m2a and m2b mutants, carrying the C-to-T and TC-to-GT mutations respectively, were constructed (Fig. 5A) and used to infect *N. benthamiana* plants. As shown in Fig. 5B, in the infiltrated leaves both m2a and m2b were able to replicate. However, m2a replicated to visibly lower levels than m2b (Fig. 5B, top two panels, lanes 9-12). We then examined the systemic leaves through 6 wpi. Unlike m2-infected plants, the m2a- and m2b-infected plants developed clearly visible systemic symptoms (Fig. 5C). Consistently, the TYLCV-specific PCR products obtained from systemic leaves of m2a- and m2b-infected plants reached high levels (Fig. 5D, lanes 14-24).

**Figure 5.**
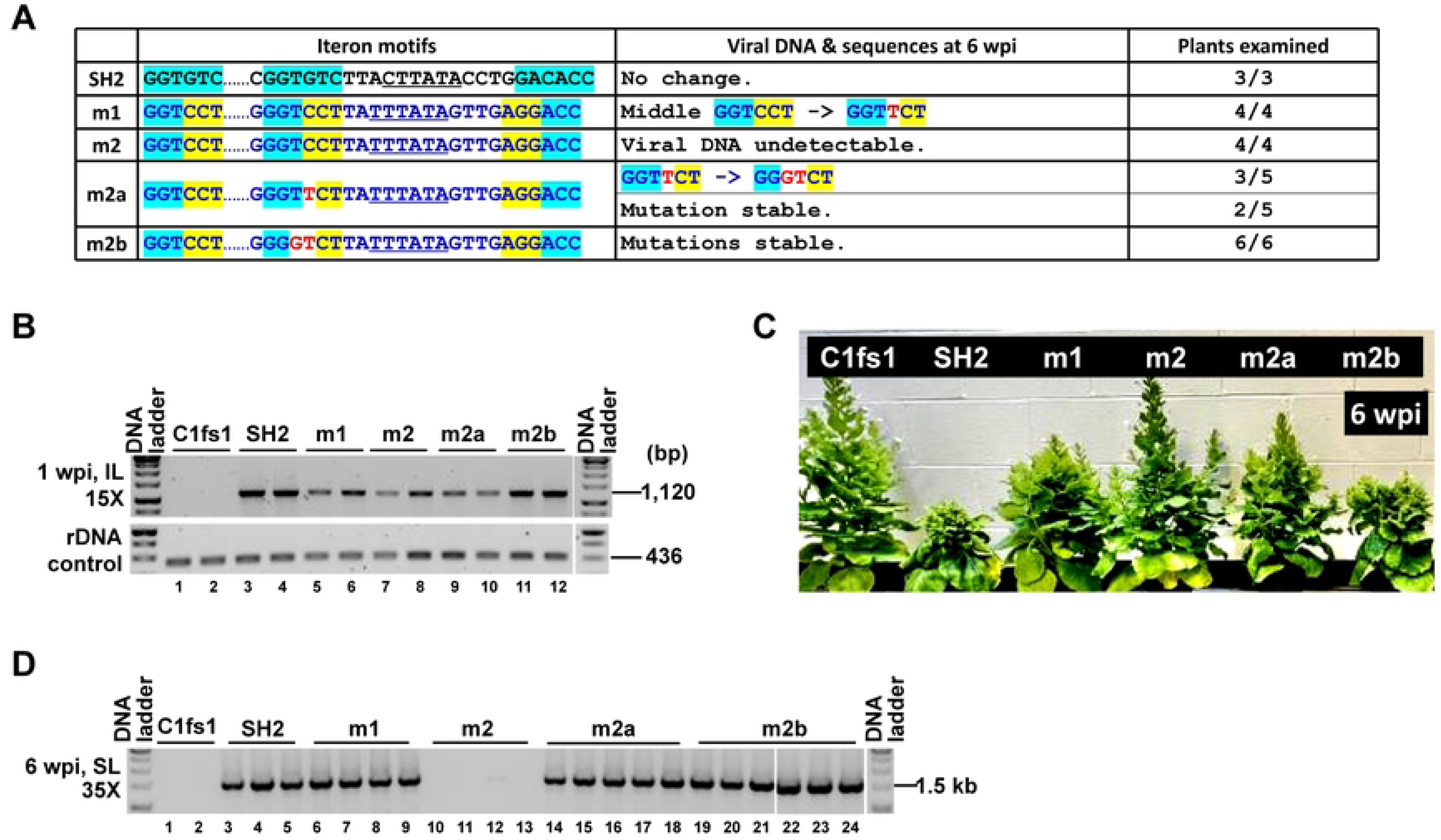
*De novo* mutations identified in m1 and m2 descendants enhance m2 symptoms. **A**. Iteron portion sequences of new m2-based mutants incorporating the *de novo* mutations; and sequences of their descendants at 6 wpi. **B**. Relative accumulation levels of SH2, m1, m2, m2a, and m2b in IL at 1 wpi. **C**. Symptoms of infected plants at 6 wpi. **D**. Accumulation levels of viral DNA in SL at 6 wpi as measured with 35 cycles of PCR. Note the absence of viral DNA in three of the m2-infected plants, and the extremely low accumulation in the fourth.

Since the m1 mutant was also included in the current experiment as one of controls, we analyzed the sequences of PCR products obtained from m1-infected samples once again. As shown in Fig. 5A, 4 of 4 analyzed sequences contained the C-to-T mutation that was identified repeatedly (Figs. 3A, 4A, 5A). Interestingly, while the C-to-T mutation was very stable in the m1 background (e.g. m1a in Fig. 4), it was less so in the m2 background (m2a). By 6 wpi, the m2a mutant acquired an additional T-to-G change at the adjacent position in 3 of the 5 plants, rendering the descendants identical to m2b (Fig. 5A). Consistent with the increased infectivity attributable to the TC-to-GT mutations in m2 (but not m1) descendants, the m2b mutant caused more pronounced symptoms than m2a in infected plants, and the mutations remained stable for at least 9 weeks. Collectively these data indicate that the *de novo* mutations occurring in m2 genome were necessary and sufficient for the new mutants to gain robust systemic infections.

### The Y35-borne middle motif represses TYLCV systemic infections even without the proximal and distal motifs, and this repression is always relieved by the TC-to-GT mutations

Given that all 3 iteron motifs in m2 were of Y35 origin, we next examined if the proximal and distal motifs also contributed to the systemic movement failure of the m2 mutant. To this end, we generated a series of mutants that altered the proximal or distal motif alone or in combination, with or without the TC-to-GT change in the middle motif. As summarized in Fig. 6A, the failure to spread systemically persisted even when one or both flanking motifs were mutated to entirely unrelated sequences (m2g, m2h, m2g-h, m2-g2-h). Note all of them replicated in ILs to levels similar to m2 (Fig. 6B, lanes 8 & 9 show m2g as a representative example). Strikingly, the TC-to-GT change in the middle motif alone was enough to restore systemic infections to these mutants (m2b-g, m2b-h, m2b-g-h, m2b-g2-h). More importantly, these 4 TC-to-GT-containing mutants all caused symptomatic infections highly similar to m2b (Fig 6C and D, compare m2b, m2g, and m2bg). Thus, the flanking motifs are unnecessary for either the middle-iteron-mediated repression or its relief by the TC-to-GT mutations.

**Figure 6.**
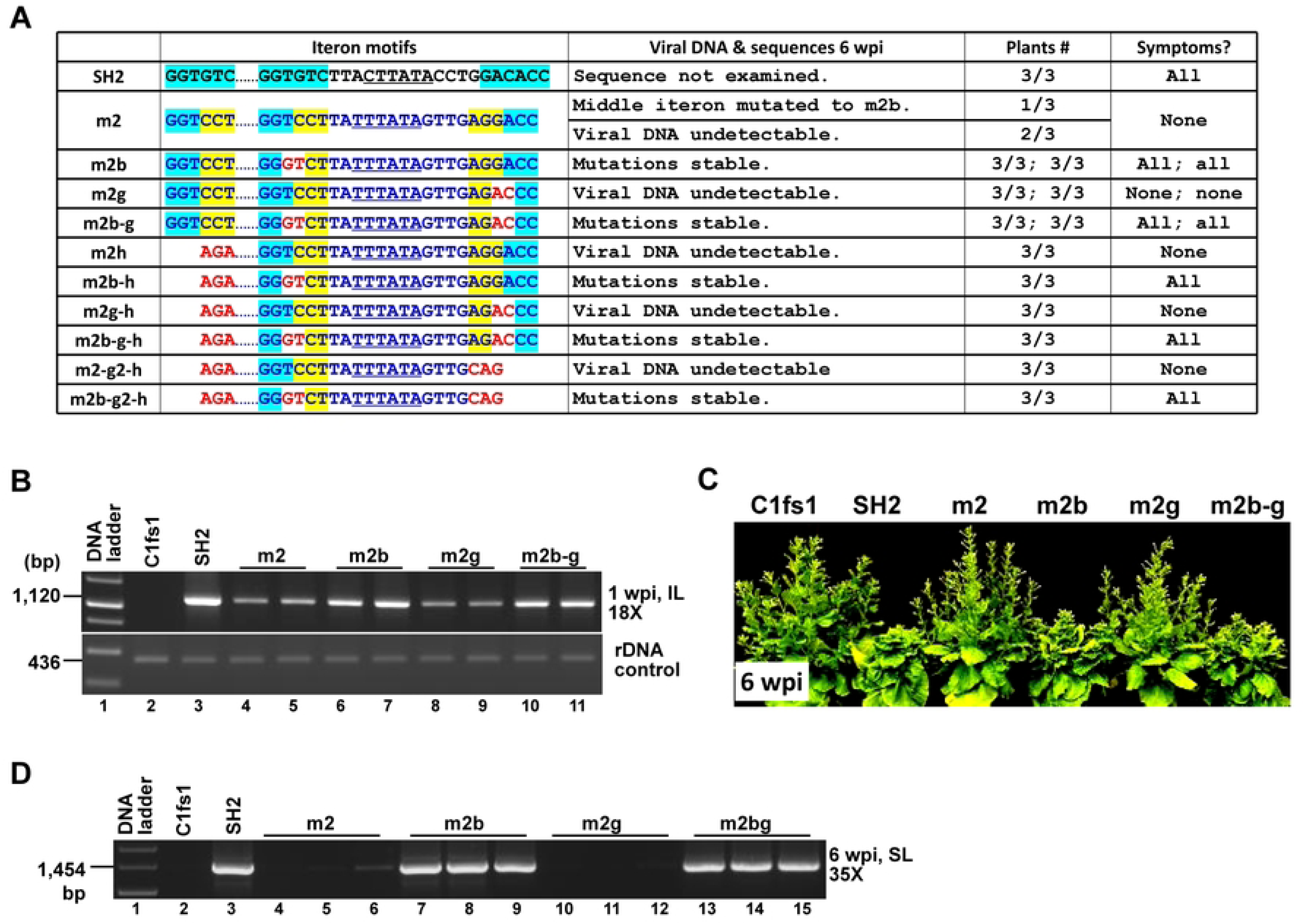
The TC-to-GT mutations of m2b are sufficient to rescue viral replication even if one or both of the flanking iteron motifs (of Y35 origin) are eliminated. **A**. Iteron portion sequences of the mutants, their absence/presence in 6 wpi SL, stability of the mutations, and SL symptoms. **B**. Semi-quantitative PCR assessing the viral DNA levels in IL for a selected set of mutants (m2g, m2b-g). **C**. Symptoms of representative mutants at 6 wpi. **D**. Detection of viral DNA in SL with PCR (35 cycles) at 6 wpi.

### Systemic infection of m2 mutant is variably bolstered by other mutations at the 3^rd^ and 4^th^ positions of middle iteron

So far we showed that the TC-to-GT mutations at 3^rd^ and 4^th^ positions of m2 middle iteron caused the resulting m2b mutant to infect *N. benthamiana* plants systemically, leading to clear symptoms indistinguishable from wildtype SH2. The intriguing puzzle was that the exact sequence of m2b middle iteron differed from that of SH2 or Y35 (GG<u>GT</u>CT in m2b as opposed to GGTGTC in SH2, and GGTCCT in Y35). We thus wondered whether substituting the TC doublet within GGTCCT – the Y35-borne m2 middle motif – with two random nts would be enough to restore systemic infection to m2. To resolve this question, we generated three new mutants – m2c, m2d, and m2e. As shown in Fig. 7A & B, m2c and m2d changed the TC doublet to AG and CA, respectively. The m2e mutant altered the last 3 nt from CCT to GTC so that this motif was now identical to that of SH2 (GGTGTC), though the rest of the iteron-encompassing section was still of Y35 origin. Furthermore, since the proximal and distal iterons were deemed inconsequential in m2b variants, we also tested three additional mutants in which m2c, m2d, and m2e changes were combined with those of m2g2 and m2h, yielding mutants m2c-g2-h, m2d-g2-h, and m2e-g2-h (Fig. 7A & B).

**Figure 7.**
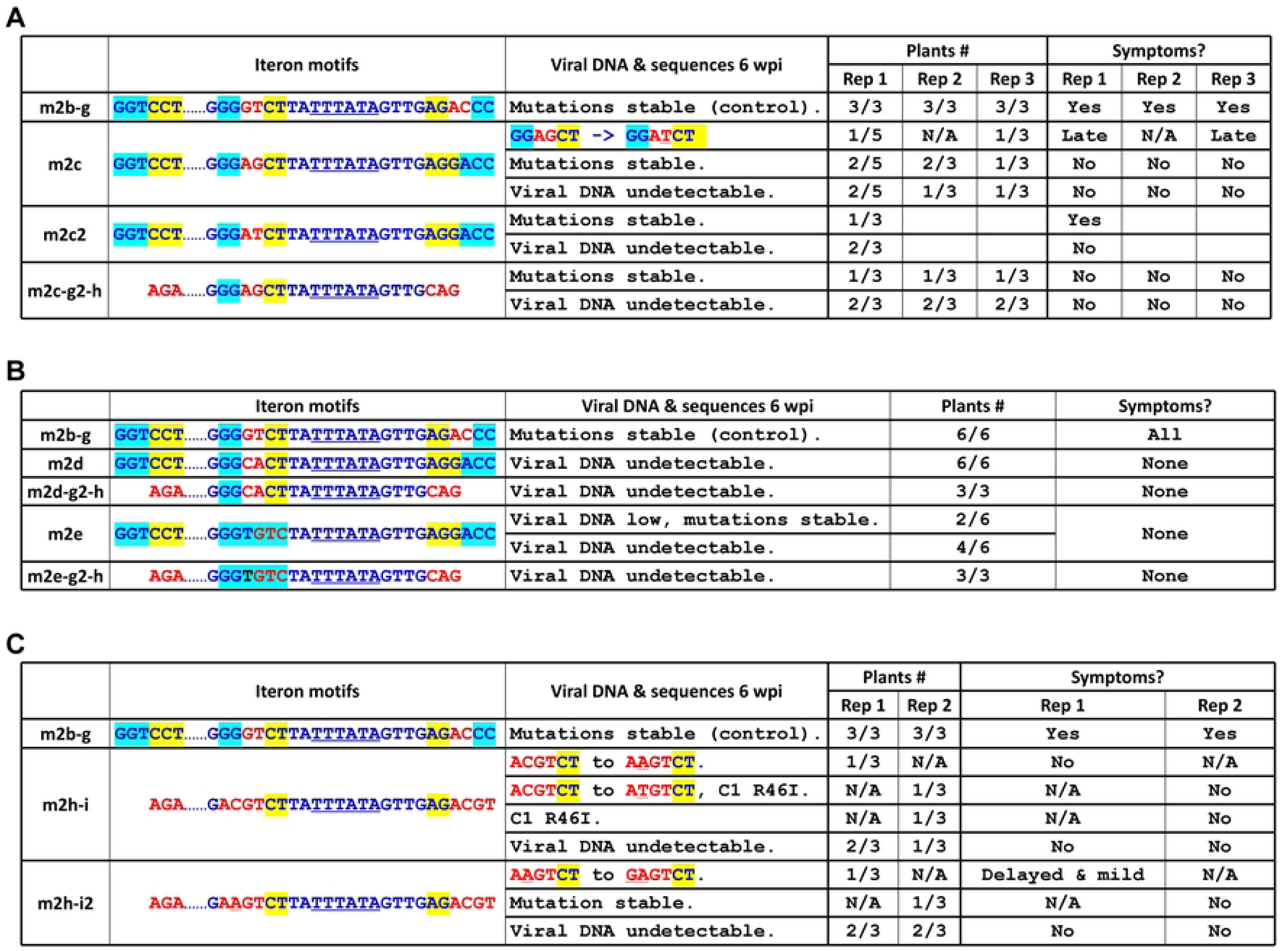
*In planta* fate of more mutants altering the two central nt of the Y35-borne middle iteron, with or without additional manipulations of proximal and distal iterons. **A**. Infection outcomes of m2c, m2c-g2-h, m2c2. **B**. Infection outcomes of m2d, m2d-g2-h, m2e, and m2e-g2-. **C**. infection outcomes of m2h-i and m2h-i2.

The m2c mutant replicated in local agro-infiltrated leaves. When the upper young leaves were assessed at 6 wpi with PCR, viral DNA was detected at in 3 of 5, 2 of 3, and 2 of 3 plants in 3 independent repeat experiments (Fig. 7A). Notably, a *de novo* G-to-T mutation at the 4^th^ position was recovered from one plant in 1^st^ trial, and another in 3^rd^ trial, changing the middle iteron motif from GGA<u>G</u>CT to GGA<u>T</u>CT (Fig. 7A). More interestingly, In both 1^st^ and 3^rd^ trials the plants from which the *de novo* mutation was recovered showed delayed symptoms that emerged at 4-5 wpi, whereas all plants containing the original m2c mutants were asymptomatic.

To further assess the impact of this *de novo* mutation, we created and tested the m2c2 mutant harboring this mutation (Fig. 7A). Indeed, the m2c2 mutant caused systemic symptoms in 1 of 3 infected plants. Importantly, the symptoms emerged at almost the same time as plants inoculated with the m2b-g control (Fig. 7A). Furthermore, the m2c2 mutations were stable in this plant, whereas the two symptomless plants were virus-free. Thus, the 4^th^ position of middle iteron motif converged to a T residue in plants receiving m1, m2, and m2c mutants, even though the respective starting residues were different (C in m1 and m2, G in m2c). By contrast, the 3^rd^ position residue could be a T (m1 progeny), G (m2 progeny), or A (m2c2 progeny). These results indicate that for symptomatic infections to occur, it was not necessary for the middle iteron to acquire a definitive sequence identity. Finally, the m2c-g2-h mutant, in which both the proximal and distal iteron motifs (of Y35 origin) were perturbed with mutations, could still be detected in unaltered form in one of three plants in three independent repeats, though none of the mutant-containing plants showed any symptoms (Fig. 7A).

By contrast, the TC-to-CA mutations of m2d mutant, with or without the flanking Y35 iterons, led to complete failure of systemic infections (Fig. 7B). Curiously, the m2e mutant, in which the SH2 middle iteron motif had been restored, albeit in the middle of Y35 context, accumulated very low levels of viral DNA in just 2 of the 5 inoculated plants by 6 wpi (Fig. 7B. The 6^th^ plant died of injury before reaching 6 wpi), with none of the plants showing any symptoms. Furthermore, eliminating the two flanking Y35 iterons led the resulting mutant (m2e-g2-h, Fig. 7B) to be completely absent from systemic leaves. The m2e and m2e-g2-h results indicated that restoring the middle iteron alone to SH2 sequence was insufficient for the virus to regain symptomatic infections. These results contrasted with those of m2b and m2c2, where departure of the middle motif from the SH2 and Y35 consensuses with merely two nt changes was enough to restore symptomatic infections. Overall they argue against a critical role of unique sequence identity for iterons. Rather, they support the argument that these *de novo* mutations primarily enabled viral escape from repression conferred by Y35 iterons in the absence of Y35 Rep.

### GG doublet at 1^st^ and 2^nd^ positions of m2b middle iteron is not essential for viral systemic spread, but probably needed for symptom manifestation

The TC-to-GT change at 3^rd^ and 4^th^ positions led to an altered motif (GG<u>GT</u>CT) that differed from Y35 (GGTCCT) and SH2 (GGTGTC). Nevertheless, all 3 motifs still shared the GG doublet at the 1^st^ and 2^nd^ positions. We next tested whether this GG doublet was needed for systemic infections, by mutating them to AC. More specifically, the m2h-i mutant changed the proximal, middle, and distal motifs to AGA, ACGTCT, and AGACGT, respectively (Fig. 1C, m2h-i). As a result, none of the 3 iterons retained the sequences of SH2 (GGTGTC/GACACC) or Y35 (GGTCCT/AGGACC), although the flanking non-iteron sequences were of Y35 origin (Fig. 7C, m2h-i). None of the 6 plants inoculated in two separate attempts showed any systemic symptoms. However, viral DNA were detected in one plant in the 1^st^ trial, two the 2^nd^ trial (Fig. 7C). Interestingly, all three viral DNA samples incurred *de novo* mutations within the middle motif and/or Rep N-terminus (Fig. 7C). In one plant, the ACGTCT was changed to A<u>A</u>GTCT. In another plant, it was changed to A<u>T</u>GTCT, plus an aa change in TYLCV Rep – position 46 arginine (R) was changed to isoleucine (I). Interestingly, the R46I change was also independently detected in another plant (Fig. 7C). Thus, the m2h-i mutant appeared to enrich compensatory changes in the Rep protein. Together these findings suggested that (i) the GG doublet was not always required for viral systemic spread, but probably played a crucial role in more robust infections that led to visible symptoms; (ii) none of the *de novo* mutations restored the GG doublet or even just one G. Rather, they appear to give rise to alternative sequence motifs, and possibly affording novel binding specificity for a co-evolved Rep.

To test whether some of the *de novo* mutations incurred in m2h-i descendants enhanced viral infectivity, we introduced one of the mutations, ACGTCT to A<u>A</u>GTCT, into m2h-i to obtain m2h-i2. The m2h-i2 mutant was still weak, detectable in one of the 3 plants in each of two independent experiments. Interestingly, another *de novo* mutation was detected in the 6-nt motif in one plant, further changing A<u>A</u>GTCT to <u>GA</u>GTCT, thus restoring one G at the 1^st^ position. These results suggest a stepwise evolutionary trajectory toward the restoration of at least one of the two Gs.

### Putative iteron-borne, intra-molecular DNA secondary structures do not predict the systemic infection differences of the mutants

Results described above demonstrated that while the Y35-borne middle iteron must incur *de novo* mutations to regain systemic infections, the exact nature of the new nt exhibited certain preference that defy easy interpretation. We hence considered the possibility that the 3 iteron motifs might fold into intra-molecular DNA secondary structures, which could then be differentially perturbed by the mutations introduced (Suppl Fig. 2). We thus used the mFold algorithm (http://www.unafold.org/mfold/applications/dna-folding-form.php) to predict potential DNA secondary structures in the iteron-encompassing sequences of SH2 (m3), Y35 (m1, m2), m1a/m2a, m1f, m1b/m2b, m2c, m2d, and m2e. As shown in Suppl Fig. 2, the m1 and m2 iterons (of Y35 origin) were indeed predicted to fold into a relative stable structure (ΔG = −5.96 kca/mol). However, this structure was weakened by m1a, m2a, m2b, m2b, m2e mutations to similar extents, hence could not explain their differences in symptom severities. Furthermore, despite folding into a much weaker structure, the m1f mutant symptoms were indistinguishable from that of m1a. Finally, the m2b-g2-h mutant, with both flanking motifs deleted, was not expected to fold into stable secondary structures, yet still elicited visible symptoms (Fig. 6A). Thus, the potential secondary structures the iterons could assume in single-stranded genomes do not provide satisfactory explanations for the observed infectivity differences.

## Discussion

### Sequence identity of iteron motifs is not essential for TYLCV replication

The critical importance of iterons with specific nucleotide sequences in geminivirus replication was first reported years ago (11,13,14). These earlier studies found that iterons with specific sequence motifs were required for replication of the bipartite tomato golden mosaic virus (TGMV) DNA-B segment, which depended DNA-A for the replication protein AL1 (11,26). Similar sequence specificity requirements for monopartite geminiviruses such as TYLCV and TbCSV were inferred from the fact these viruses likewise harbor reiterated short motifs upstream of the Rep coding region (7,9,27). However, whether iteron motifs of monopartite geminiviruses are essential for replication has not been carefully examined. To the best of our knowledge, Xu and colleagues (10) were the first to address this question. They found that mutants of the TbCSV isolate Y35 with any two of the three iterons deleted still infected plants systemically, but a mutant with all three iterons deleted was non-infectious (10).

The current study built on these earlier findings and established that none of the iteron motifs was absolutely required for TYLCV (isolate SH2) replication in initially infected cells. Compared with a mutant of SH2 containing a loss-of-function mutation in Rep (C1fs1), all of our iteron mutants, ranging from m2 in which all three of the SH2 iterons were replaced by their Y35 counterparts, to m2h-i in which all three of them were mutated to even more distinct sequences, were able to replicate to varying levels in agro-inoculated primary leaves. Moreover, some of the mutants, such as m1, m2, m2c, m2c-g2-h, and m2h-i, were detectable in systemic leaves and incur *de novo* mutations. Yet others, such as m1a/b, m2a/b, m2b-g, m2b-h, m2b-g2-h, and m2c2, elicited clear symptoms despite of extensive perturbation of some or all iteron motifs. Together our results indicate that specific iteron sequences are not essential for TYLCV replication in infected cells.

### Heterologous iterons absent of a matching Rep repress TYLCV replication

An important revelation of our study is that iteron motifs by themselves most likely repress replication of the cognate geminivirus. To explain our reasoning, let us first note that Rep proteins are thought to contain two separate domains mediating iteron-binding (9). The first domain corresponds to N-terminal 8-10 aa, whereas the second maps to aa 66-75 of SH2 Rep, or 64-73 of Y35 Rep (Suppl Fig. 1). The m1 mutant acquired from Y35, along with the iteron-containing non-coding region, the N-terminal iteron-binding domain of Y35 Rep. Thus, a low level of iteron-Rep binding may still occur between Y35 iterons and the matching Rep N-terminus. While the m1 mutant reproducibly elicited systemic symptoms, the descendant viruses always contained the G-to-T *de novo* mutation at 4^th^ position of middle iteron, converting the motif from GGTCCT to GGT<u>T</u>CT. Put differently, a single mutation within a 105-nt heterologous sequence was enough to rescue robust infections. Therefore, the imported Y35 iterons must have repressed viral replication, necessitating the C-to-T change to escape the repression.

This interpretation is further supported by the fact that the m2 mutant, in which the imported Y35 iterons had to co-exist with the non-matching SH2 Rep, failed systemic infections in most plants. Instead, only the descendants that acquired one or two *de novo* mutations within the middle iteron (GGTCCT to GGT<u>T</u>CT or GG<u>GT</u>CT, m2a and m2b) regained symptomatic infections. Therefore, compared to m1, the further absence of Y35 Rep N-terminus in m2 caused m2 to be more debilitated, making it necessary to acquire two mutations to relieve the repression imposed by Y35 iterons. We hasten to note that the original m1 and m2 mutants were both able to replicate at modest levels in cells they first entered, and we surmise that such low-level replication was necessary for *de novo* mutations to emerge and proliferate.

To reiterate, even though the m1 and m2 mutants differed from the wildtype SH2 by sequence stretches of 105 and 60 nt, respectively, acquiring one and two of *de novo* mutations within the imported sequences was enough to restore symptomatic infections. Especially given that both the proximal and distal motifs still retained Y35 sequences in the infection-generated m1 and m2 descendants, it is unlikely that a stimulative role could be regained by altering just one of two nt. It is much more conceivable to foresee such minor changes relieving a certain repressive activity conferred by iterons and/or nearby sequences.

### The *de novo* mutations likely overcome the iteron-mediated repression by evading certain replication-blocking features of iterons and/or nearby DNA sequence(s)

If iterons by themselves repress replication, it can be inferred that in wildtype TYLCV infections, the cognate Rep acted to defeat this repression through iteron-Rep binding, thereby facilitating genome replication. How could iteron-mediated repression be defeated when a matching Rep was unavailable? Results with our mutants illustrated that in the absence of a matching Rep, iteron-mediated repression could be overcome by incurring one or two *de novo* mutations within the middle iteron motif. It is worth repeating that the new middle motifs (GGT<u>T</u>CT, GG<u>GT</u>CT, GG<u>AT</u>CT) created through *de novo* mutations did not match that of wildtype SH2 (GGTCTC), thus unlikely to have rescued the original iteron-Rep binding. Further disputing the involvement of a specific sequence was that one of the new middle motifs, GG<u>GT</u>CT, potentiated symptomatic infections even when both proximal and distal iterons (of Y35 origin) were eliminated (the m2b-g2-h mutant). Conversely, restoring the middle motif to that of SH2 in the Y35 context was insufficient to rescue symptomatic infections (the m2e mutant), even after the Y35 proximal and distal iterons were both eliminated (the m2e-g2-h mutant). Thus, instead of recreating a new sequence motif to suit SH2 Rep, the *de novo* mutations must have bolstered systemic infections by overcoming repression conferred by the Y35 iterons.

How do Y35 iteron motifs repress TYLCV replication? To resolve this puzzle, we considered the possibility that iteron motifs, or other sequence motifs that either overlap with iterons or tightly linked to them, serve as the binding sites for host-encoded transcription factors (TFs). The rationale for this idea is that recruitment of TFs to iterons, which are closely linked to the TATA-box of the promoter driving Rep mRNA transcription, likely maximizes Rep production immediately after viral genomes enter host cell nuclei (28). Nevertheless, heavy TF attachment to this iteron-containing genome section likely blocks the same section from being accessed by Rep and other replication-related proteins. Consistent with this idea, iteron motifs are invariably found upstream of the Rep coding sequence (27). Furthermore, their repetition for 3-4 times within a relatively short stretch closely resembled the arrangement of TF binding sites in the promoters or enhancers of many cellular genes (29–31). For example, the 6-nt auxin-responsive promoter element (AuxRE, TGTCTC) were frequently found multiple times in promoters of auxin-responsive genes, in the form of tandem and/or inverted repeats. Indeed, synthetic promoters with repeated AuxRE motifs were shown to be much more potent than native ones, and were frequently used to screen TFs that bind to the motif (29,32).

Further supporting this idea was the identification of diverse TF binding sites within genomes of various geminiviruses, many of which shown to mediate transcriptional enhancement of downstream genes (33–35). Particularly relevant to our discussion is the G-box motif (CACGTG) found in the TGMV DNA-A segment, within the 5’ non-coding region of the TGMV-encoded Rep (AL1). G-box motifs are binding sites for TFs of two large families – the basic helix-loop-helix (bHLH) family and basic leucine zipper (bZIP) family (11,36,37). The G-box motif in TGMV DNA-A is separated from iterons by a 28-nt region that contained the TATA box (38). This G-box-plus-TATA-box-plus-iterons region was verified as a functional promoter driving strong transcription of a reporter gene (35,37,38). Strikingly, this promoter activity was all but abolished when the G-box motif was mutated, suggesting the involvement of G-box-binding TFs in AL1 mRNA transcription (35). Even more interestingly, this promoter activity was potently repressed by the TGMV AL1 protein through sequence-specific iteron-AL1 binding (37). These observations strongly suggest that TF-binding to G-box motif activates AL1 mRNA transcription, whereas Rep-binding to iteron motifs represses the same transcription activity, likely by peeling TFs off the promoter through competitive binding (35).

The non-coding region of SH2 genome encompassing iterons does not contain a G-box motif. However, it does contain two consecutive repeats of a different motif, AATTCAAA, 36 nt upstream of iterons. AATTCAAA is a near-perfect match for the binding sites of three Arabidopsis TFs, TSO1, TCX2, and TCX3 (https://jaspar.elixir.no/search?q=&collection=CORE&tax_group=plants). TSO1, TCX2, and TCX3 are closely related TFs of the CPP (cysteine-rich polycomb-like) family found to play crucial roles in Arabidopsis reproduction by coordinating the cell fate determination in both male and female reproductive organs, and controlling stem cell division (39,40). Separately, the middle iteron motif of SH2, if extended by 2 nt upstream, has the sequence of *TC*GGTGTC, or GACACC*GA* on the complementary strand. Within these 8 nt, TCGGTG/CACCGA is highly enriched in the binding sites of multiple ethylene response factors (ERFs), including ERF011, 019, 037, 038, and 043. Conversely, the Y35 middle iteron along with 2 upstream nt would have the sequence of *TG*GGTCCT/AGGACC*CA*, containing TGGGTCC/GGACCCA, a motif enriched in the binding sites of multiple TCP TFs, including Arabidopsis TCP7, 9, 21, and 22 (https://jaspar.elixir.no/search?q=&collection=CORE&tax_group=plants). TCPs are TFs implicated in diverse processes such as floral symmetry control, branching, lateral organ development, as well as defense responses (41). Notably, the TGGGTCC motif would have been changed by m2b mutations into TGGG<u>GT</u>C, possibly compromising the binding by the same set of TCPs.

### The <u>a</u>ntagonistic <u>b</u>inding of iterons by <u>T</u>Fs and <u>R</u>eps (ABITR) model

Together these observations prompted us to propose the ABITR model (ABITR) (Fig. 8). ABITR postulates that iterons are tightly linked to, or overlap with, binding sites of certain host-encoded TFs. Geminiviruses evolve TF-binding sites to lure TFs to the Rep promoter, ensuring rapid production of Rep protein. Yet occupation of Rep promoter by TFs also makes the occupied genome copies recalcitrant to replication proteins. This in turn select for iteron motifs nearby, facilitating iteron-Rep interaction that peels off TFs, availing the TF-free genome copies for replication (Fig. 8). Based on this model, iterons are kept in replication-repressing states through TF occupation. Conversely, *de novo* mutations within the TF-binding sites/iterons, such as those in m2b, likely relieved the repression by lessening TF binding to Rep promoters, permitting replication initiation despite the absence of specific iteron-Rep binding.

**Figure 8.**
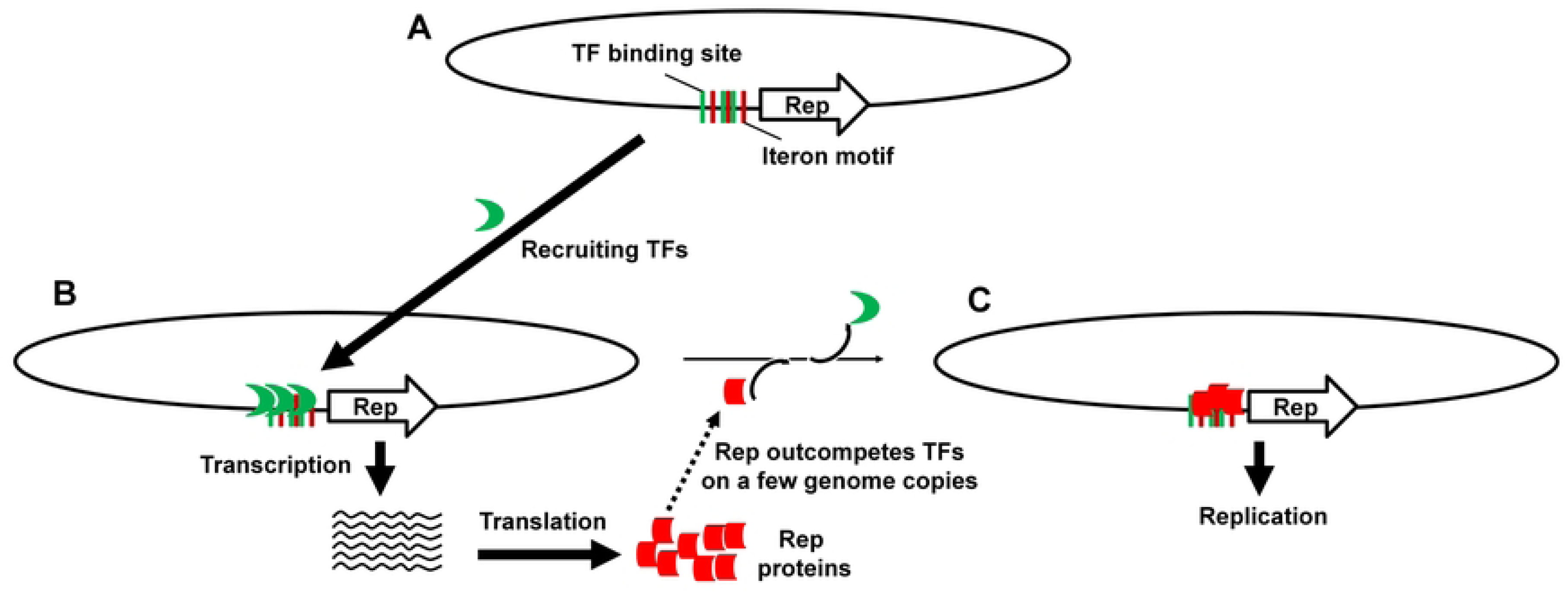
The schematic depiction of antagonistic binding of iterons by TFs and Reps (ABITR). **A**. Multiple TYLCV genome copies co-entering the same cell nucleus enlist the iteron-proximal TF binding sites to recruit TFs. **B**. Recruitment of TFs to multiple genome copies activates synchronous Rep mRNA transcription, ensuring rapid accumulation of Rep protein. **C**. At an appropriate concentration threshold, Rep proteins compete with TFs for iteron motifs and nearby TF binding sites, succeeding in routing a few genome copies per cell to replication.

### Antagonistic iteron-binding by TFs and Reps likely bottlenecks geminivirus replication and drives iteron motif diversification

We previously reported that TYLCV replication was intracellularly bottlenecked so that among dozens of genome copies that entered a cell, no more than three could initiated replication (16). Such stringent intracellular reproductive bottlenecking is critical for sustaining viral viability as it enables swift purging of defective genome copies harboring lethal and deleterious errors (42–44). However, it is not yet known how geminiviruses like TYLCV establish such intracellular reproductive bottlenecks. The ABITR arrangement, if verified through additional future research, could constitute an early stage of intracellular bottlenecking. This is because, upon cellular entry, the iteron-encompassing TF-binding sites on Rep promoter should be immediately available for binding with host-encoded TFs. Such promoter-TF binding should be sufficiently stable, hence preventing the subsequently synthesized Rep proteins from de-repressing more than a few genome copies. For the bottlenecks to sustain, there is abundant evidence showing Rep over-accumulation later during geminivirus cellular infection blocks, rather than facilitates, viral replication. Indeed, Rep overexpression has been widely used to engineer resistance to geminiviruses (45–49). In short, intracellular reproductive bottlenecking of geminiviruses is likely first established with the help of host-encoded TFs, and later refortified by over-accumulated Rep proteins.

Conversely, novel TF-binding specificity could emerge if a viral genome copy incurs mutation(s) allowing it to escape TF-imposed reproductive bottlenecking, hence replicating to dominance in cells containing Rep-supplying sister copies. However, such bottleneck-evading cheater copies are bound to accumulate excessive numbers of lethal errors through unconstrained replication. The descendant lineages consisting of such cheaters, upon entering new cells, are expected to either cease replication, or acquire additional mutations that together recreate binding sites for a different set of TF(s). This then set in motion the evolution for new iteron motifs, and also new binding specificities in Rep. Such “Red-Queen” race would explain the rapid diversification of geminivirus iterons.

To summarize, our extensive investigations of TYLCV iterons yielded the surprising revelation that specific iteron sequence motifs are not required for viral replication. Rather, they act as repressors of TYLCV replication, and their interaction with cognate Rep protein serves to overcome this repressive activity. Moreover, absent of a cognate Rep, the repressive activity of iterons can also be overcome by incurring relatively few (one or two) *de novo* mutations, thus unveiling a different sequence specificity constraint in addition to iteron-Rep binding. Based on these results, we propose the ABITR model postulating that sequence motifs in the close vicinity of iterons likely evolve sequence specificity matching certain host-encoded TFs, thereby recruiting TFs to enhance Rep mRNA transcription and Rep protein production. Once Rep accumulates to a certain concentration threshold, it probably expels TFs from a limited number of viral genome copies through competitive iteron binding, availing these genome copies to bottlenecked replication. Testing predictions of this new ABITR model through future research will likely unveil novel targets for more effective management of crop diseases caused by geminiviruses.

## Materials and Methods

### Constructs

The original TYLCV infectious clone (isolate SH2. The Genbank accession number is AM282874.1) was kindly provided by Dr. Xueping Zhou of China Institute of Plant Protection (15). The full-length, double-stranded form of TYLCV genome, plus a 50-bp duplication at the 5’ end, and another 50-bp duplication at the 3’ end, was subcloned into pAI101, an *E.coli*-*A. tumefaciens* shuttle vector modified from pCambia1300 in our lab (50,51), leading to a new TYLCV infectious clone we call LM5. To create LM5-G and LM5-R, two KpnI sites were introduced into the V1 gene, at positions 307/308 and 881/882 (numbering relative to the full length genome sequence), respectively. The coding sequences of uvGFP and mCherry were then PCR amplified and cloned between the KpnI site using the NEBuilder kit (New England Biolabs). The sequences of LM5, LM5-G and LM5-R encompassing the entire TYLCV DNA (and its modified forms) were verified with Sanger sequencing. Several previously generated constructs (e.g. P35S-p19) were also used (52–54).

Most of the mutants were generated by producing the mutation-containing PCR fragments through overlapping PCR (sequences of the primers used are available upon request). The resulting PCR fragments were cloned into the LM5 infectious clone digested with SgsI and Bsp119I. A few mutants were generated with custom-synthesized, mutation-containing DNA fragments (TWIST Biosciences).

### Agrobacterium infiltration (agro-infiltration)

All DNA constructs destined for testing in *N. benthamiana* plants were transformed into electrocompetent *A. tumefaciens* strain C58C1 via electroporation using the AGR setting on the Bio-Rad Micropulser Electroporator. Briefly, 1 µl of the plasmid DNA (100-250 ng) was mixed with 40 µl of agro cells and maintained on ice until electroporation. After electroporation, 900 µl of SOB media was added and the suspension was incubated at 28 °C for one hour. Selection was carried out on solid Terrific Broth (TB) media containing rifampicin, gentamycin, and kanamycin. Successful introduction of the plasmid was confirmed using colony PCR. A single colony confirmed to have the desired plasmid was used to inoculate 3 ml TB liquid media with the same antibiotics, and incubated overnight at 28 °C. The culture was diluted 1:100 with fresh TB liquid media and incubated under the same conditions for another night. The second culture was centrifuged at 4,000 rpm for 20 min, and resuspended in agroinfiltration buffer (10 mM MgCl2, 10 mM MES, and 100 µM acetosyringone). All suspensions were diluted to OD600 = 0.5 and incubated at room temperature for 3 hours. *Agrobacterium* suspensions were then mixed and introduced into leaves of young *N. bethamiana* plants via a small wound, using a needleless syringe.

### Confocal microscopy

Four days after agro-infiltration, leaf discs were collected from the plants. Confocal microscopy was performed at the Molecular and Cellular Imaging Center (MCIC), the Ohio Agricultural Research and Development Center, using a Leica DMI6000 laser confocal scanning microscope. To detect GFP and mCherry fluorescence, sequential excitation at 488 nm and 587 nm was provided by argon and helium-neon 543 lasers, respectively.

### Assessing viral replication in inoculated leaves with semi-quantitative PCR

DNA was extracted from agro-infiltrated leaves at 1 wpi using the Quick-DNA Plant Kit manufactured by Zymo Research. The DNA samples were adjusted to the same concentration prior to PCR. A pair of primers, SH2-2157F and SH2-495R (Fig. 1A, light blue arrows above the genome drawing. Primer sequences available upon request), were used to generate a replication-dependent fragment of 1,120 bp. The non-replicating mutant C1fs1, which contained a 1-nt insertion in Rep coding region causing Rep reading frame to shift, was included in these experiments serving as a negative control. The PCR was carried out for 15 or 18 cycles to avoid low-level circular forms generated independently of replication. In most experiments a ribosomal DNA fragment of 436 bp was PCR-amplified in parallel to serve as controls for similar amounts of plant DNA.

### Detecting *de novo* mutations via Sanger sequencing

The systemic leaf DNA samples were obtained with the Quick-DNA Plant kit (Zymo Research) for some experiments, and the REDExtract-N-Amp Plant Kit (Sigma-Aldrich) for others. For the purpose of Sanger sequencing, a pair of PCR primers that amplified the entire Rep coding region plus the intergenic region between Rep and V2 were used (SH2-1475F and SH2-196R; sequences available upon request). PCR was usually repeated for 35 cycles, and the amplified fragments were sequenced by the OSU Genomics Core Facility.

## Acknowledgements

We thank Dr. Xueping Zhou of China Institute for Plant Protection for generously sharing the SH2 infectious clone; and the USDA-ARS Maize, Wheat, and Soybean Viruses Group for generous equipment sharing. Members of the Qu lab are greatly appreciated for discussions and technical assistance. This study is supported in part by the NSF grant 1758912. RFR is supported in part by a scholarship from China Scholarship Council, and the Postgraduate Scientific Research Innovation Project of Hunan Province (QL20210118).

## Supporting Information (SI)

The following SI files are available online: Supplementary Figure 1 and 2.

